# Reliability of Mismatch Negativity Event-Related Potentials in a Multisite, Traveling Subjects Study

**DOI:** 10.1101/768408

**Authors:** Brian J. Roach, Ricardo E. Carrión, Holly K. Hamilton, Peter Bachman, Aysenil Belger, Erica Duncan, Jason Johannesen, Gregory A. Light, Margaret Niznikiewicz, Jean Addington, Carrie E. Bearden, Kristin S. Cadenhead, Tyrone D. Cannon, Barbara A. Cornblatt, Thomas H. McGlashan, Diana O. Perkins, Larry Seidman, Ming Tsuang, Elaine F. Walker, Scott W. Woods, Daniel H. Mathalon

**Author notes:** Address for correspondence: Daniel H. Mathalon, Ph.D., M.D., San Francisco VA Healthcare System/ Psychiatry Service (116D), 4150 Clement Street, San Francisco, CA 94121. Indicates shared first authorship (Brian J. Roach, Ricardo E. Carrión).

## Abstract

**Objective:** Mismatch negativity (MMN) is an auditory event-related potential (ERP) used to study schizophrenia and psychosis risk. MMN reliability from a multisite, traveling subjects study was compared using different ERP referencing, averaging, and scoring techniques.

**Methods:** Reliability of frequency, duration, and double (frequency+duration) MMN was determined from eight traveling subjects, tested on two occasions at eight EEG laboratory sites. Deviant-specific variance components were estimated for MMN peak amplitude and latency measures using different ERP processing methods. Generalizability (G) coefficients were calculated using two-facet (site, occasion), fully-crossed models and single-facet (occasion) models within each laboratory to assess MMN reliability.

**Results:** G-coefficients calculated from two-facet models indicated fair (0.4<G<=0.6) duration MMN reliability at electrode Fz, but poor (G<0.4) double and frequency MMN reliability. Single-facet G-coefficients averaged across laboratory resulted in improved reliability (G>0.5). Reliability of MMN amplitude was greater than latency, and reliability with mastoid referencing significantly outperformed nose-referencing.

**Conclusions:** EEG preprocessing methods have a significant impact on the reliability of MMN amplitude. Within site MMN reliability can be excellent, consistent with prior single site studies.

**Significance:** With standardized data collection and ERP processing, MMN can be reliably obtained in multisite studies, providing larger samples sizes within rare patient groups.

## Introduction

Mismatch negativity (MMN) is an event-related potential (ERP) component that is automatically elicited by an infrequent deviant auditory stimulus that differs in pitch, duration, or another sound feature from a repetitive series of preceding “standard” stimuli. MMN is considered to reflect sensory echoic memory, since the detection of auditory deviance depends on the short-term online formation of a memory trace of the immediately preceding standard sounds in the auditory processing stream, and can be measured using either electroencephalography (EEG) or magnetoencephalography (MEG). The MMN has great potential as an ERP biomarker because of its robust sensitivity to the pathophysiology of schizophrenia [14, 39, 2] and its ability to predict transition to a psychotic disorder in individuals at clinical high-risk (CHR) [31, 4, 36]. While the test-retest reliability of MMN has been the focus of several studies witihin a single laboratory, the reliability and consistency of the MMN response across testing location must be evaluated in order to determine the suitability of this ERP component for use in multi-site, clinical trials or longitudinal studies of psychosis.

Many prior test-retest reliability studies of MMN relied upon Pearson [30, 38, 20, 40, 35, 21] or Spearman [13, 34] correlation coefficients. This approach is somewhat limited in that such coefficients only evaluate the degree to which MMN responses or ranks from two tests covary, without considering whether responses are in close agreement from one test occasion to the next. A better measure of such agreement is the intra-class correlation (ICC) coefficient [37]. Eight studies [23, 18, 22, 24, 3, 33, 9, 25] have reported ICCs of MMN. In general, these studies have found that MMN reponses are stable over time, further highlighting the potential for the component to be used a biomarker in high risk populations [26]. Regardless of the reported coefficient type, the majority of the above mentioned studies have only evaluated or compared the impact of different paradigms or sound features used to elicit the MMN. To the best of our knowledge, no study has assessed the influence of different EEG signal processing choices, such as the reference electrode or methods of artifact rejection, on the reliability of the MMN responses despite the fact that these choices differ across reports.

In multi-site studies of the reliability of functional magnetic resonance imaging (fMRI) data, generalizability (G) theory has been applied to facilitate descriptions of the different sources of variance in blood oxygen level-dependent (BOLD) signal measurements [6, 15]. G-theory applies a random effects modeling approach to partition sources of variance by calculating variance components for the effect of persons as well as other measurement factors, or facets (e.g., study site, testing occasion), and their interactions. The variance components can then be used to calculate generalizability or dependability (G- or D-) coefficients. The G-coefficient is relevant when relative measurements or differences between subjects are of interest (e.g., a 3 µV difference between MMN responses from a patient and control subject) while the D-coefficient is relevant when absolute measurements are of interest (e.g., a patient has a −3µV MMN response). The identification of critical facets and estimation of associated variance components is considered a G-study in the G-theory framework. While the G-study and estimated variance components are sufficient to calculate both G- and D-coefficients, the theory separately labels optimization of reliability coefficients for future studies or data collection procedures as a decision study (or D-study). In the D-study, estimated variance components are used to determine how many measurements are required to produce a sufficiently high G- or D-coefficient, when averaging across facets such as test item or occasion.

The present study focuses on a two facet, fully-crossed traveling subjects G-study of MMN in the North American Prodrome Longitudinal Study (NAPLS), which is a multi-site research consortium studying the mechanisms and predictors associated with psychosis onset. The two facets were Site (NAPLS geographic location) and Occasion (test and retest day). The fully-crossed design of this G-study indicates that all persons had MMN measured on two occasions at each study site. Variance is partitioned into the main random effects of Person, Site, and Occasion, their two-way interactions, and a final term corresponding to the three-way interaction plus error. In the case of a multi-site G-study, such as the current study, the Person X Site variance component would be part of the denominator of both G- and D-coefficients, but the Site variance component would only be included in the denominator of the D-coefficient. Thus, the D-coefficient will be less than the G-coefficient if the Site, Occasion and Site X Occasion variance components are non-zero. As these three components approach zero, the D- and G-coefficients converge. The present G-study design is identical to previously published physiology [7], fMRI [15, 27] and structural MRI [8] reports from the NAPLS consortium because the same traveling subjects underwent EEG and MRI (for more details please see [1]) assessments on each test day. The main goals were to (1) Quantify variance components and associated G-coefficients representing the single site, single session reliability of MMN measured with different scoring methods, averaging approaches, and reference electrodes; (2) Compare reliability of MMN using average mastoid versus nose references:, and (3) Compare reliability of MMN using two different averaging methods. Lastly, since previous MMN reliability studies were conducted at a single laboratory site, individual laboratory site G-coefficients were also calculated separately for each of the 8 NAPLS 2 sites and a “home” site model to allow for qualitative comparisons within the NAPLS consortium and the extant literature.

## Methods

### Participants

#### Traveling Subjects Sample

One healthy participant was recruited from each of the eight NAPLS2 sites. EEG data were collected on two consecutive test days at each site, starting at each participant’s home site followed by a pseudo-random travel order to all other sites. The average number of days between the first test occasion at each site was 7.3 (SD=7.6), and particpants completed all 16 EEG sessions in 29 to 80 days. Participants were between 19 and 31 years old (mean=27.74, SD=3.99), and there were an equal number of males and females. All participants provided written informed consent and the protocol was approved by the Institutional Review Boards at each of the NAPLS sites.

### Equipment

#### EEG

All participating data collection sites used BioSemi (www.biosemi.com) EEG acquisition systems. Half (UCLA, Harvard, UNC, Yale) of these systems were equipped to record 64 channels of EEG, and half (Emory, Hillside, UCSD, Calgary) were equipped to record 32 channels.

#### Stimulus Presentation

All sites used Dell Optiplex Desktop computers to run the MMN task (described below). These systems were configured to meet or exceed the minimum hardware requirements recommended at the time of study launch (2009) by the stimulus presentation software provider, neurobehavioral systems (www.neurobs.com), with special attention paid to video and sound cards. LCD monitors connected to a 512MB ATI Radeon PCIe video card by VGA cables were used at each site. Auditory stimuli were delivered via ER1 Etymotic insert earphones connected to a SoundBlaster X-Fi Xtreme Gamer PCI card. Subject responses were recorded with a Cedrus RB-830.

### Paradigms

#### Hearing Test

Auditory stimuli were presented through ER1 Etymotic insert earphones using Presentation software. Prior to the MMN task, hearing levels were also assessed using the same stimulus presentation software and hardware employed in the MMN task. The hearing thresholds for three pure tones (500, 633, and 1000 Hz) were detected separately for each ear at the beginning of every session. This was accomplished by playing 50 ms duration tones of each frequency in each ear, manipulating the “attenuation” parameter within the software. The attenuation value was set between 0 (no attenuation) and 1 (total attenuation). Starting with an attenuation value of 1, the value was decreased in 0.05 increments until the subject indicated that she had heard a tone in the target ear by pressing a left or right response button corresponding to the ear in which the tone was detected. A 0.05 step in attenuation is theoretically equivalent to 5 dB, but in practice the actual change in dB depends on the auditory stimulus delivery device and its frequency response function. Before starting the test, subjects were told to respond to the tones played through the right or left ear insert, and that the tones would never be played in both ears at the same time. After the subject’s first response to a specific tone, the attenuation value was increased by 0.2 or set to 1, whichever was less, and the process was repeated until the subject detected a tone from each frequency in each ear four times.

#### MMN Paradigm

Auditory stimuli delivery consisted of 85% standard tones presented for 50 ms at 633 Hz, 5% duration (DUR) deviants presented for 100 ms at 633 Hz, 5% frequency (FRQ) deviants presented for 50 ms at 1000 Hz, and 5% double-deviants (DBL) presented for 100 ms at 1000 Hz. A total of 1794 tones were presented over 3 separate blocks, with each block lasting approximately 5 minutes. Tones were presented with 5 ms rise and fall times and a 500 ms stimulus onset asynchrony. In an effort to reduce the effect of attention on the MMN ERPs, participants were instructed to ignore auditory stimuli while focusing on a separate distractor task. The distractor task consisted of a visual oddball paradigm that was run simultaneously with MMN, and the presentation of the visual stimuli were jittered to avoid co-occurring visual oddball and MMN ERP signals.

### EEG Collection, Preprocessing, and ERP Averaging

#### Data Acquisition

EEG was recorded at 1024 Hz using either a 32-channel or 64-channel electrode cap. Additional electrodes were placed on the face and mastoids, and an offline average mastoid reference was used for the following data analysis. In the present study, data from one subject on one test occasion were incomplete due to equipment malfunction, and required the elimination of some sections of the continuous recording where shorting of the electrodes occurred. As a result, less than half of the total trials were available for this test session from the start of data preprocessing. For one additional subject and test occasion, operator error resulted in no continuous EEG recording in one of the three test blocks. Due to the complicated study design and small sample size, both of these recordings were included in all reliability analyses.

#### Preprocessing

EEG recordings were re-referenced to average mastoids and high-pass filtered at 1 Hz before being segmented into 1000 ms epochs (−500 to 500 ms). Blinks and eye movement artifacts were recorded by electrodes placed around the eyes and were corrected for by using the ocular correction method outlined in Gratton, et al. [17]. Following baseline correction (−100 to 0 ms), outlier electrodes were interpolated within single trial epochs based on previously established criteria [28]. A spherical spline interpolation [12] was applied to any channel that was determined to be a statistical outlier (|z| > 3) on one or more of four parameters, including variance to detect additive noise, median gradient to detect high-frequency activity, amplitude range to detect pop-offs, and deviation of the mean amplitude from the common average to detect electrical drift. Epochs with amplitudes greater than ±100 µV in any of the following electrodes were rejected: AF3, AF4, F3, Fz, F4, FC1, FC2, FC5, FC6, C3, Cz, C4.

#### ERP Averaging and MMN Measurement

ERP averages for all stimulus types were determined using a sorted averaging method [32]. This method has been shown to reduce noise in the MMN waveform by averaging over the subset of trials that optimizes the estimated signal to noise ratio (eSNR) for each subject. In this data set, single-epoch root mean squared (RMS) amplitude values at each of the 12 electrodes used for artifact rejection for each trial were calculated, averaged across electrode, and sorted in ascending order for each stimulus type. The subset of sorted trials selected for ERP averaging were associated with the largest eSNR, which is the ratio of the number of trials to the variance of the amplitude values across trials. To facilitate comparison with more traditional MMN processing methods, a separate set of ERP averages were also obtained ommiting this sorted averaging step. Following averaging, ERPs for all stimulus types were low-pass filtered at 30 Hz, and then standard tone ERP waves were subtracted from deviants to obtain difference waves. As the reference electrode could influence both artifact rejection and sorted averaging trial elimination steps, nose re-referencing was done on the final waveforms to facilitate reference electrode comparisons on the exact same set of trials. MMN peak amplitude was classified as the most negative peak between 90 and 290 ms in each calculated difference wave. MMN mean amplitude ±10 ms around the peak was also quantified as an alternative measurement to peak amplitude. Finally, average amplitude in a fixed window based on grand average waveforms (90-170 ms for FRQ and DBL, 150-230 for DUR) was quantified as a third approach. Peak latencies were saved for a fourth set of generalizability analyses.

### Traveling Subject Sample Reliability Analyses

#### Variance Components and G-coefficients

The main purpose of this fully crossed, two facet (site and test occasion) G-study design is to estimate variance components. This particular design allows one to estimate 7 variance components for any given score from each person, at each site, on each test occasion. The variance components and their definitions are described in Table 1.

**Table 1.**
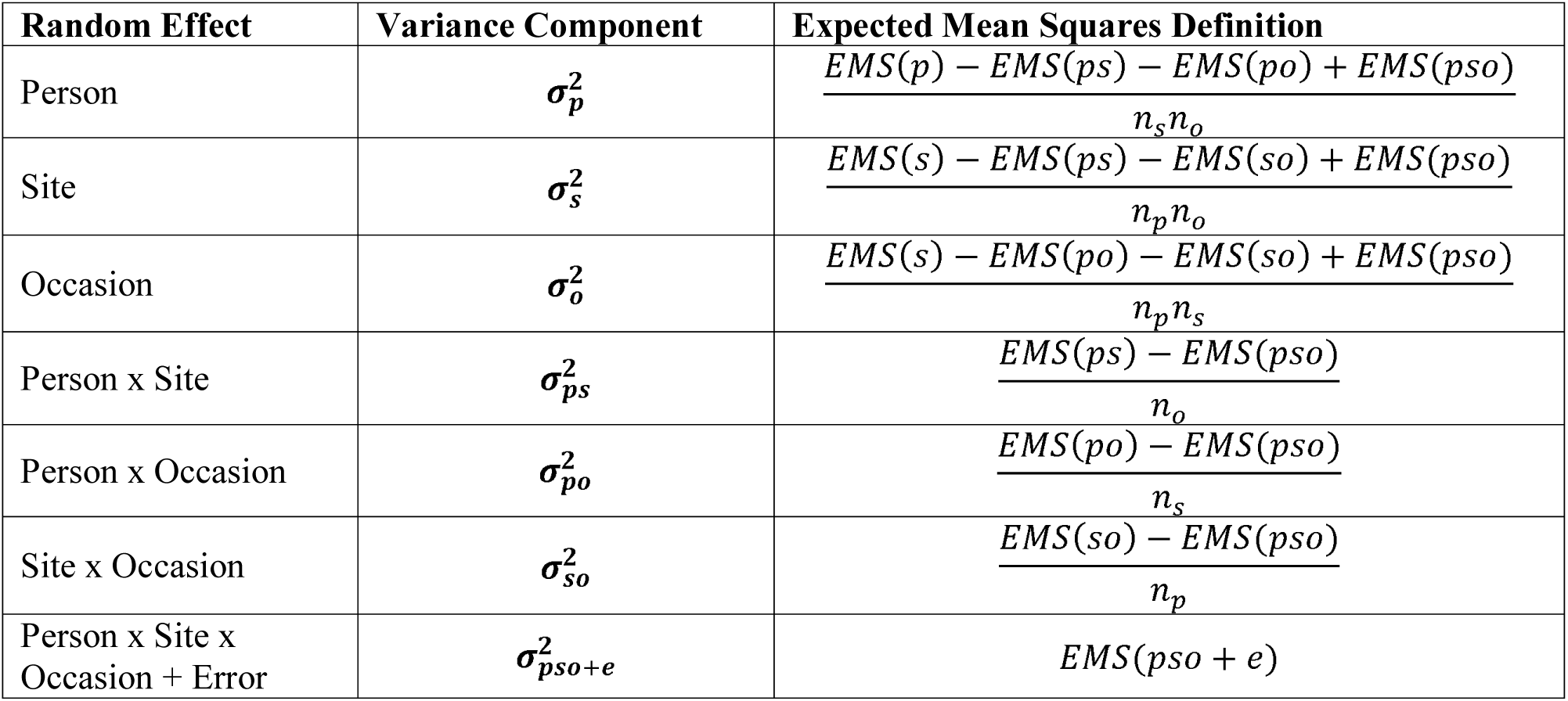
Variance Components for *p* x *s* x *o* Fully Crossed Design

Once variance components are estimated, the G-coefficient, which provides a measure of generalizability or reliability of the measured score, can be calculated as:

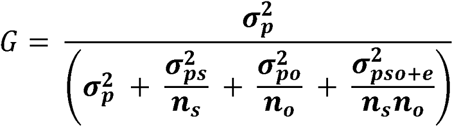

The larger NAPLS2 parent study design included EEG assessments at baseline, 12 month, and 24 month study time points. Since MMN scores from each session would be treated separately, with particular emphasis on using baseline data to predict conversion to psychosis in the parent study, the logical choice for ***n_o_*** is 1. Likewise, since all subjects would only be studied at their home site, the logical choice for ***n_s_*** is 1. Therefore, the G-coeffecient is equivalent to the intraclass correlation (ICC) as defined by Shrout and Fleiss (e.g., ICC(3,1) in [37]) when ***n_o_*** = ***n_s_*** = 1. Variance components were estimated using a restricted maximum likelihood approach implemented in Matlab [41]. Components were estimated and saved separately for each deviant type (DBL, FRQ, DUR), electrode (32 from overlapping montage), MMN measurement (peak amplitude, mean around peak, mean in fixed window, and peak latency), ERP averaging method (sorted or traditional) and reference electrode (average mastoids or nose).

While the visual oddball attention task and hearing tests are not the focus of this report, variance components were also estimated for median target reaction time (RT), overall accuracy in the oddball task, and mean hearing thresholds from the hearing test.

#### Additional Generalizability Analyses

To assess the influence of individual subjects and sites on the estimated variance components, a delete-1 jackknife approach was used to calculate 80 sets of variance components (8 delete-1 site, 8 delete-1 person, and 8*8 delete-1 site and person). In all jackknife models, both test ocassions were used to maintain the functional form of the two facet model described previously, and analyses were limited to electrode Fz both to reduce computations and because of its frequent use in MMN research.

Finally, as the most frequently reported MMN reliability studies collect data on two test occassions at one laboratory site, 9 separate sets of reduced variance components were estimated for a single facet (test occasion) crossed design within each of the eight sites, and a final model where each subject’s initial pair of test occasions from their home site were used (“home” site model). In this last model, person and site are completely confounded, so associated variance components and G coefficients must be interpreted with this limitation in mind.

Because the “home” site reliability seemed to outperform other sites, we tested its performance with two post-hoc methods. First, we counted the number of G-coefficients (across averaging methods, deviant types, and measures for mastoid referenced data) that were greater than or equal to 0.6 (i.e., “good” or better) and compared this count to the G-coefficients from (i) the 8 NAPLS sites and (ii) the 7 pairs of subsequent visits, in which participants were equally distributed across all sites much like the home site case. Additionally, the percentage of these home site G-coefficients that were maximal were tested against the null hypothesis that maximum G-coefficients would be equally distributed either (i) across the 8 NAPLS sites and home site (e.g., P(maximum G) = 1/9 or 11.11%) or (ii) across pairs of ordered visits (e.g., P(maximum G = 1/8 or 12.5%).

### Averaging Method and Reference Electrode Comparisons

As the purpose of the G study is not to test a specific hypothesis, there are no p-values associated with estimated variance components or G-coefficients. However, existing guidelines for determining practical or clinical significance of ICCs suggest that the reliability coefficient can be qualitatively categorized as follows: ICC < 0.4 is poor, 0.4 <= ICC < 0.6 is fair, 0.6<= ICC < 0.75 is good, and 0.75 <= ICC < 1 is excellent [10]. Therefore, G-coefficients were categorized using these 4 labels and subjected to tests comparing (1) traditional versus sorted averaging methods using mastoid referenced data and (2) mastoid versus nose referenced data as follows. Cochran-Mantel-Haenszel (CMH) tests were conducted to check proportional odds assumptions across deviant type (DBL, FRQ, DUR) as well as MMN measurement type (peak, mean around peak, fixed window mean, and latency). Finally, the proportion of G-coefficients that were greater in the sorted averaging approach compared to the traditional approach using mastoid reference data were determined and tested against the null hypothesis that both averaging methods performed equally well (i.e., the proportion would equal 50%). Mastoid and nose referenced data were similarly compared.

## Results

### Variance Components and G-coefficients

MMN ERP waveforms from electrode Fz are plotted in Figure 1. There is clear similarity between waveforms at each site and on each test occasion up until about 200 ms, followed by greater variability in the 200-400 ms range.

**Figure 1.**
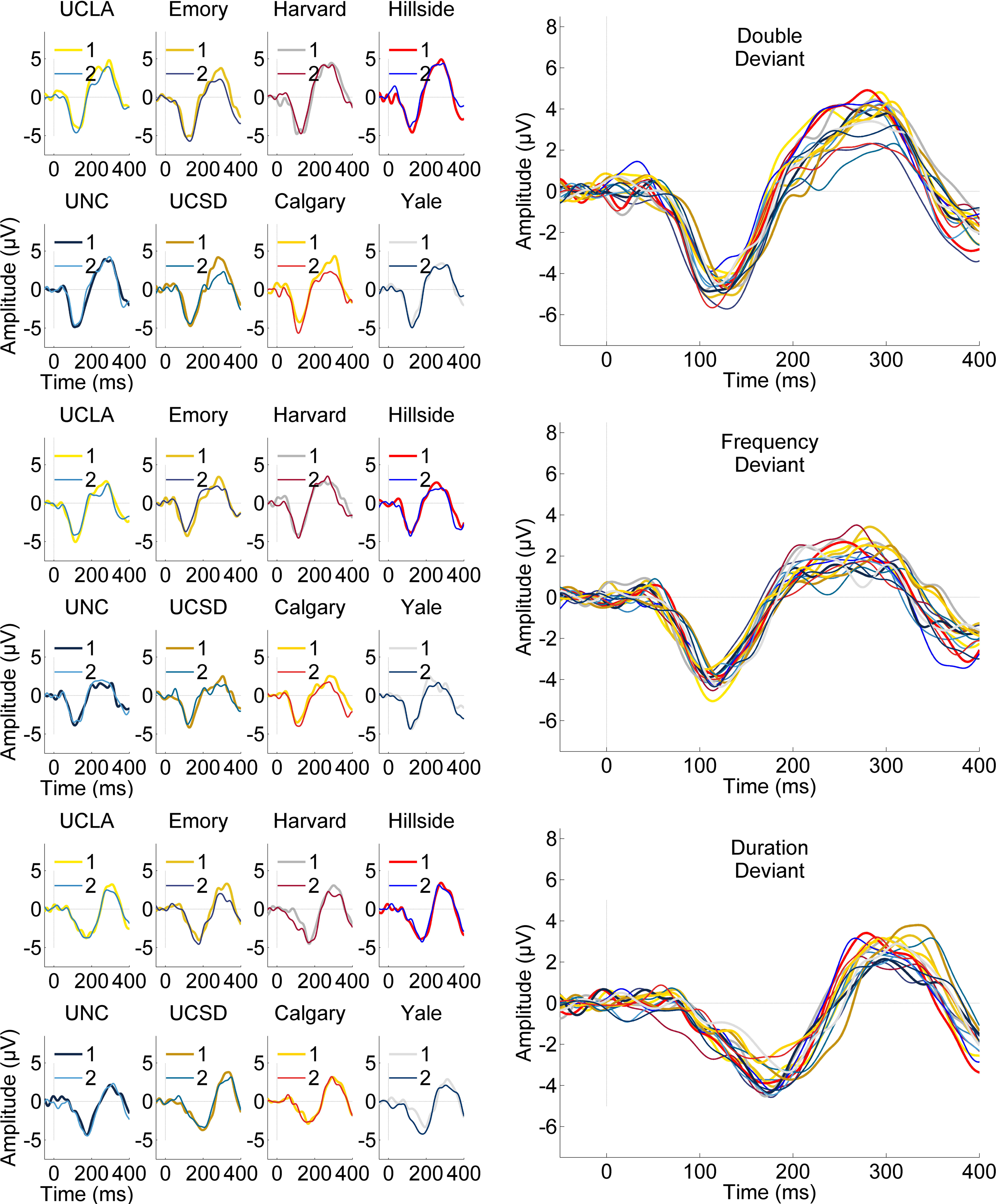
Site and session-specific grand average mismatch negativity (MMN) deviant minus standard tone difference waveforms are plotted for the Double (Frequency plus Duration) Deviant (Top), Frequency Deviant (Middle), and Duration Deviant (Bottom) from electrode Fz. Grand Average MMN waveforms for each NAPLS laboratory site are plotted separately on the right-hand side for the first (1) and second (2) test occasion. All 16 of these average waveforms are overlaid for each deviant type on the left-hand side. Time, in milliseconds (ms) from tone onset is plotted on the x-axis, and amplitude, in microVolts (µV), is plotted on the y-axis.

A similar set of waveforms was produced for the traditional averaging approach (Figure S1). Accepted trial numbers for the two averaging approaches are included in Table 2. Sorted averaging resulted in the rejection of between 6% - 7% of the trials for each trial type on average. In some sessions, sorted averaging rejected no additional trials for the deviants (range: 0 – 18 trials), while at least 10 trials were rejected from the standards in each session (range: 10 – 218 trials).

**Table 2.**
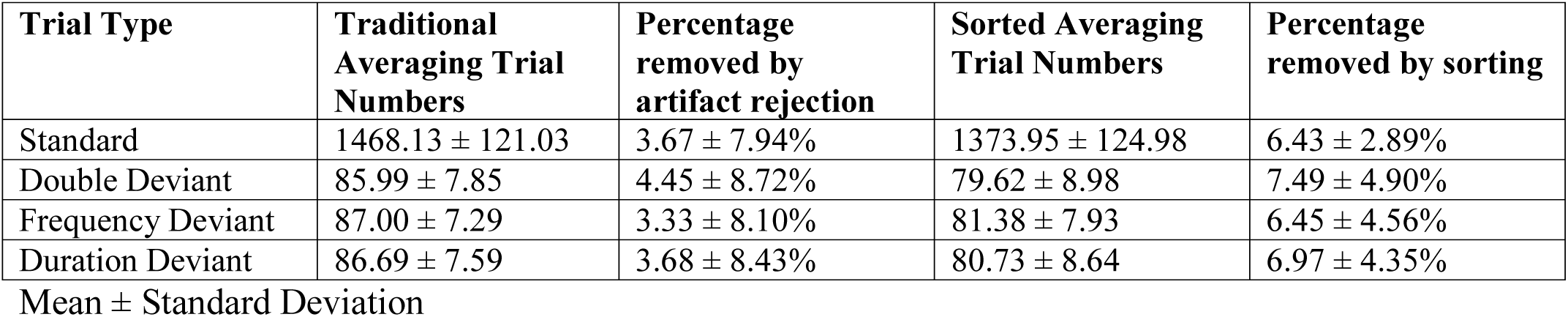
Trial Numbers

Figure 2 shows the grand average MMN waveforms from electrode Fz across all 128 sessions along with G-coefficient waveforms. While the sorted and traditional grand average ERPs are almost identical, the G-coefficient waveforms are less consistent between methods with all samples falling well below 0.4, indicating poor reliability. G-coefficients for each electrode, deviant type, reference, avereaging approach, and measure are included in Supplementary Table 1 (Table S1). The peak and mean around the peak G-coefficients for Fz are greater (0.275-0.487) than any G-coefficient in the waveforms, indicating that latency jitter in the MMN may contribute to poor sample-wise reliability in the waveforms.

**Figure 2.**
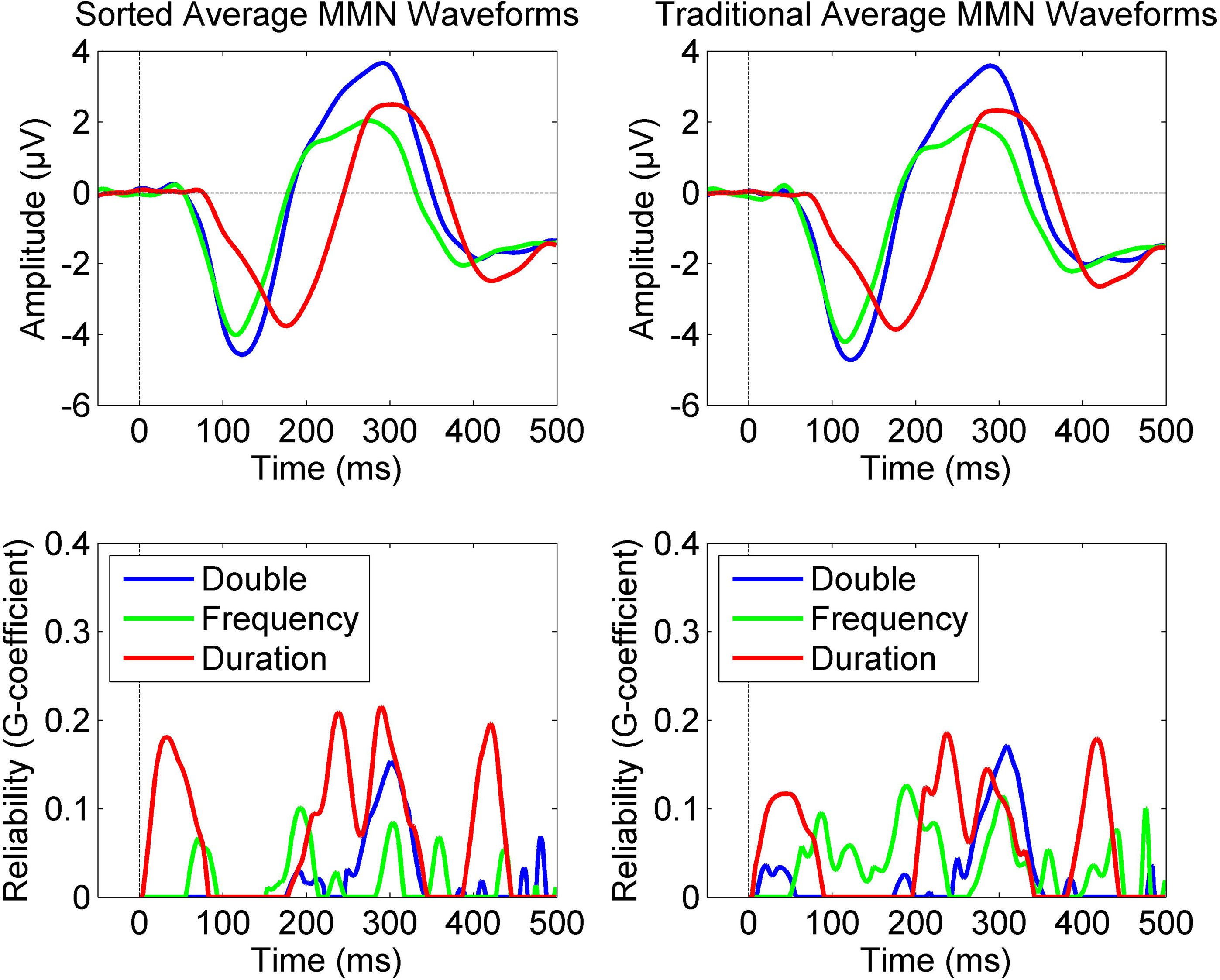
In the top row, grand average mismatch negativity (MMN) deviant minus standard tone difference waveforms are plotted for the Double (Frequency plus Duration) Deviant (blue), Frequency Deviant (green), and Duration Deviant (red) from electrode Fz. Sorted averaging (left) and traditional averaging (right) MMN waveforms are very similar in these grand averages across all 128 test sessions. Test-retest reliability waveforms are plotted separately for each deviant type and averaging method in the bottom row. These g-coefficient waveforms are derived from a two-facet (site and test occasion) fully-crossed generalizability analysis and demonstrate that the reliability (or generalizability) of the MMN waveform at any given time sample is relatively poor, indicating that MMN scores should be calculated with some averaging across time samples or peak-picking approach.

Table 3 lists the proportion of variance for each of the 7 components as well as G coefficients averaged across the 6 fronto-central electrodes (F3, Fz, F4, C3, Cz, C4) and deviants. Such averaging is consistent with the group analysis approach applied to MMN data in other CHR [31] and schizophrenia [19] studies.

**Table 3.**
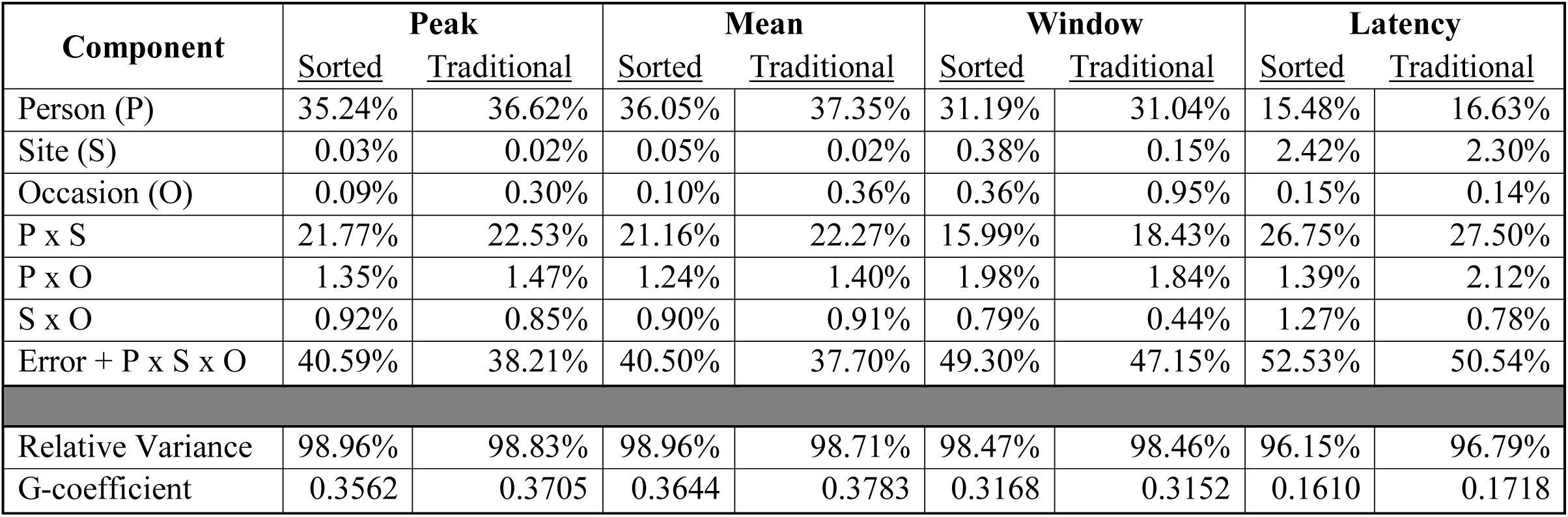
Mean proportion of variance for each Variance Component

As shown in Table 3, the two averaging approaches yielded similar estimates. This is consistent with data from Fz only as plotted in Figure 3, which shows similar percentages of variance attributed to each of the 7 variance components, when additionally separated by each deviant type. The general pattern shows that the largest variance component is error, followed by Person and then Person X Site for the three amplitude measures. For the peak latency measure, error variance still dominates, but Person X Site variance is greater than person variance. Table 4 shows the range of the proportion of variance associated with each of the 7 variance components.

**Figure 3.**
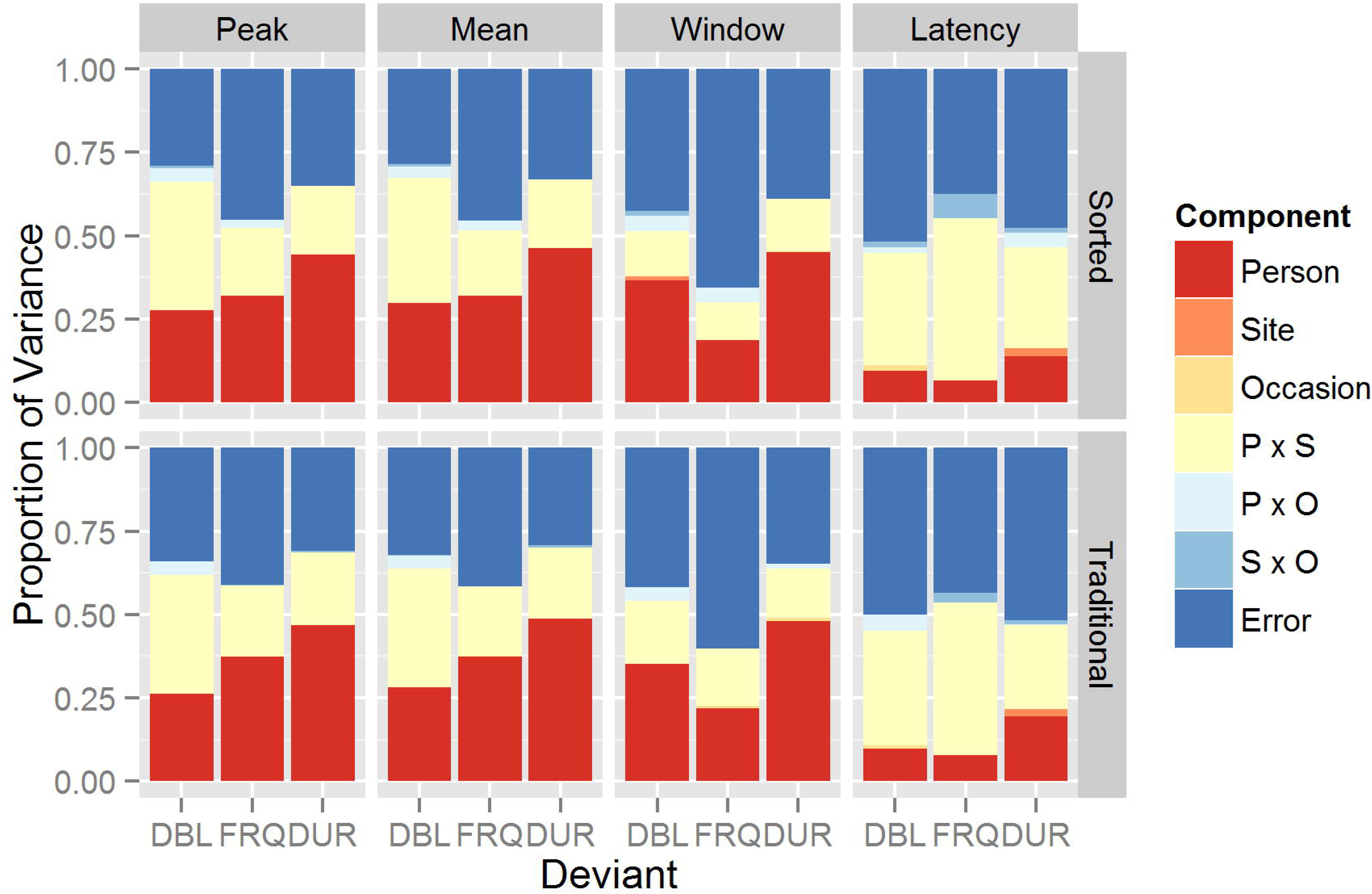
Stacked bar plots show the proportion of variance (y-axis) explained by the 7 different variance components estimated using the two-facet, fully-crossed models. The Person (red), Person x Site (yellow), and Residual Error (dark blue) variance components account for the most of the variance for mismatch negativity peak amplitude (far left), mean amplitude ± 10 milliseconds around the peak (middle left), mean amplitude in a fixed time window (middle right), and peak latency (far right) measures. Double (Frequency plus Duration; DBL), Frequency (FRQ), and Duration (DUR) Deviants are plotted separately along the x-axis from electrode Fz, and are separated by sorted averaging (top row) and traditional averaging (bottom row) methods of event-related potential calculation.

**Table 4.**
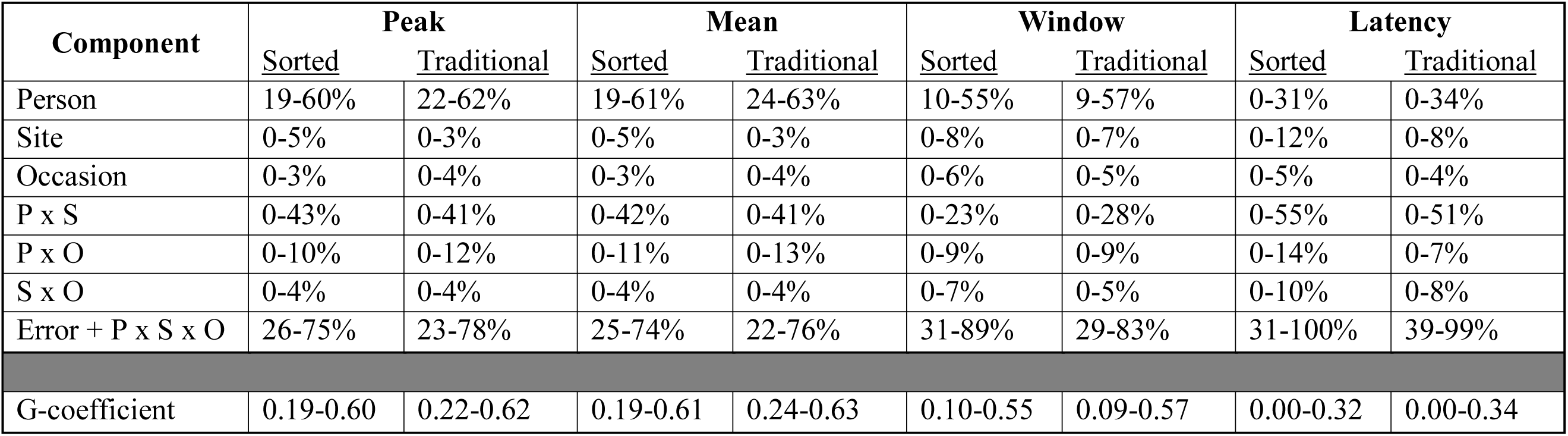
Range of proportion of variance for Variance Components in Delete-1 Analyses

Across measures and averaging methods, G-coefficients and associated person variance components fit a general pattern where DUR is greatest followed by FRQ for Peak and Mean measures or DBL for Window and Latency measures. Topographic maps on the mean amplitude in the fixed time windows and their corresponding G-coefficients for mastoid referenced, sorted average data are shown in Figure 4. Corresponding plots for traditional average data and peak amplitude are included as Figures S2-4. Much like the data from electrode Fz, the Window measure reliability follows a pattern of DUR > DBL > FRQ. All three deviant types exhibit a left-lateralized central-parietal reliability maximum, and the topographies of the scored amplitude do not perfectly match the associated G-coefficient topographies. While the reliability of DBL is improved by the Window measure and DUR reliability is unchanged relative to Peak or Mean measures, FRQ deviant reliability is reduced.

**Figure 4.**
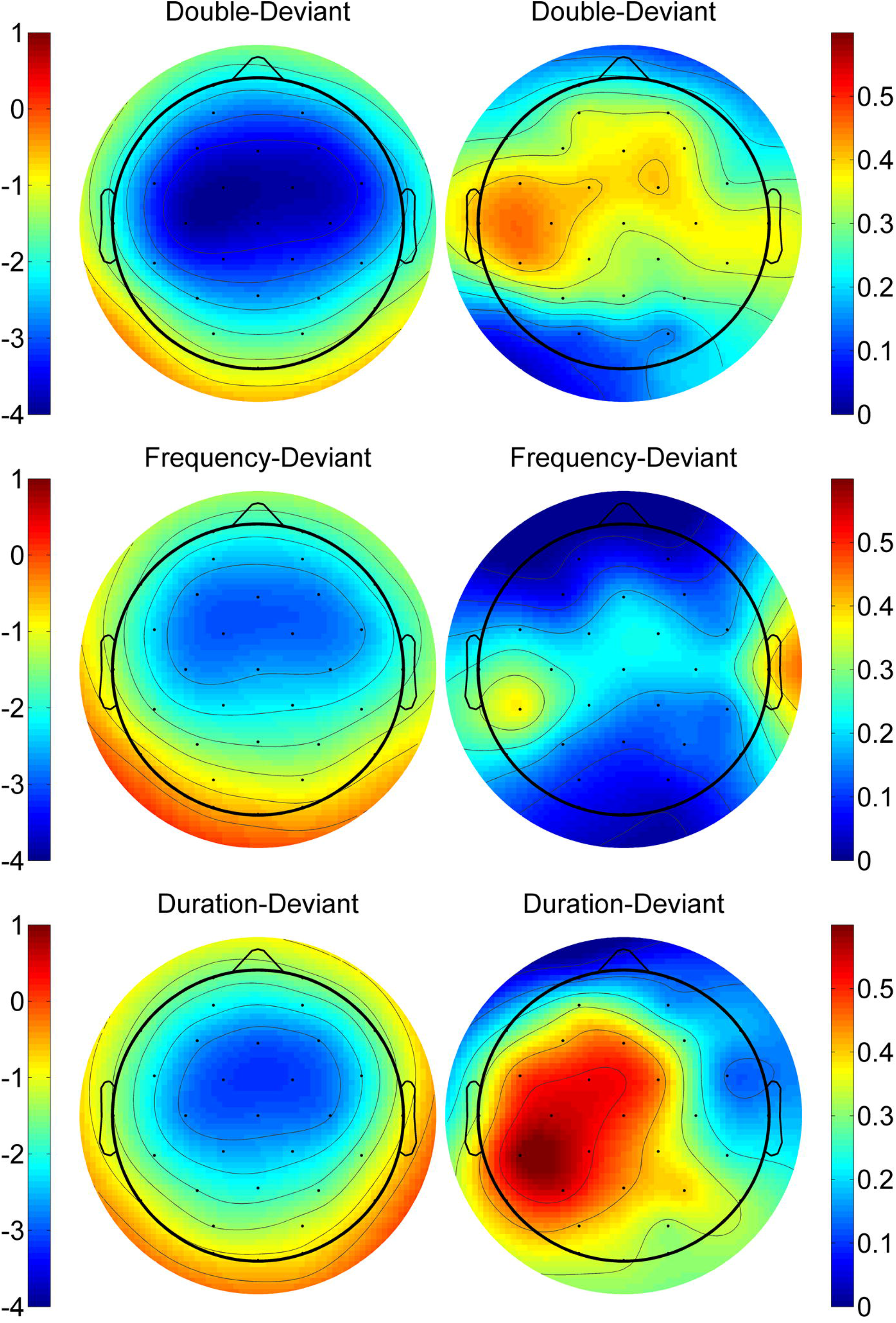
Scalp topographic maps displaying the mismatch negativity (MMN) quantified as the mean amplitude in fixed latency windows on the left-hand side, and corresponding G-coefficients for the fully crossed two-facet (site and test occasion) generalizability study on the right-hand side. Frequency deviant (middle row) and double (duration+frequency) deviant MMN is averaged across 90-170 milliseconds (ms) while Duration deviant (bottom row) MMN is averaged across 150-230 ms.

Overall oddball accuracy reliability was poor (G = 0.2327), given the overall lack in variability and ceiling level performance across most sessions (Median Accuracy = 100%, interquartile range: 99.77% - 100%). Median target RT (Median RT = 364.7, inter-quartile range: 353.3 to 393.5ms) reliability was good (G = 0.6505); the only non-zero variance components in addition to Person and the Error terms were Site and Person X Site (2.45 and 11.92% variance explained, respectively). Taken together, these measures indicate that subjects performed the visual distraction task consistently across sessions. The mean hearing level reliability was fair (G = 0.4961) with a larger proportion of variance attributed to Site (12.01%) than most other measures studied. Person X Site and Site X Occasion variance components were also non-zero (7.77 and 2.08% variance explained, respectively).

### Additional Generalizability Analyses

Figure 5 shows the mean G-coefficients and ranges from the delete-1 jackknife estimates at electrode Fz, which follows the ordering pattern for the three deviant types described previously.

**Figure 5.**
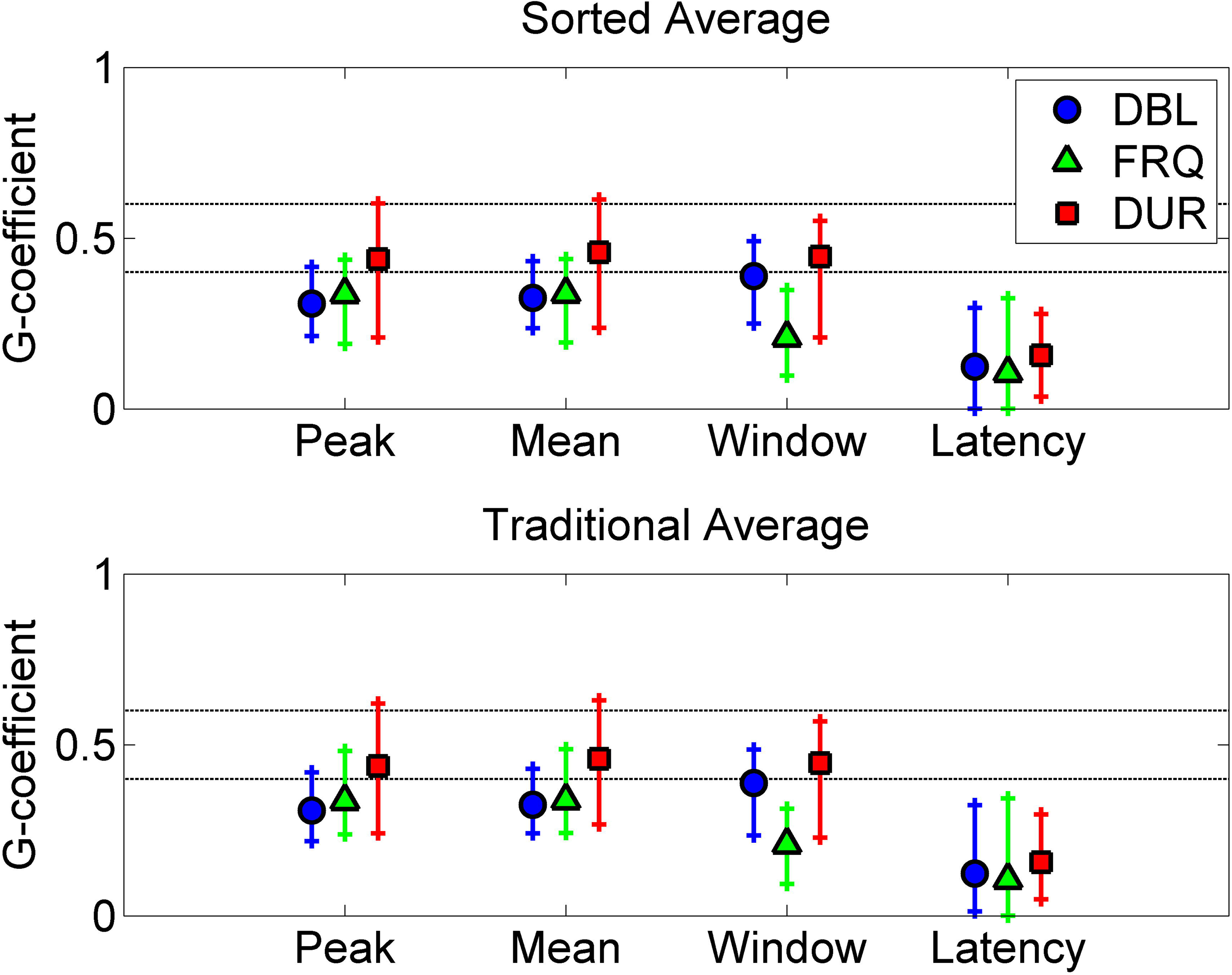
The average G-coefficients and associated ranges for the fully crossed two-facet (site and test occasion) generalizability studies across delete-1 (Site, Subject, or Site and Subject) are plotted for electrode Fz based on either sorted averaging (top) or traditional averaging (bottom) event-related potential calculation. In both cases, mismatch negativity measurement scoring approaches are plotted along the x-axis separately for double-deviant (DBL, blue circles), frequency-deviant (FRQ, green triangles), and duration-deviant (DUR, red squares). Horizontal dashed lines indicate the Fair (G = 0.4) and Good (G = 0.6) reliability levels.

When broken down by deviant type, it is clear that the mean G-coefficient and associated ranges are both greater for all amplitude measures of the DUR MMN compared to the other deviants (see Figure 5). A similar pattern can be seen when the G-coefficients are calculated separately on a per site basis (see Figure 6) using a reduced, single-facet (Occasion) model.

**Figure 6.**
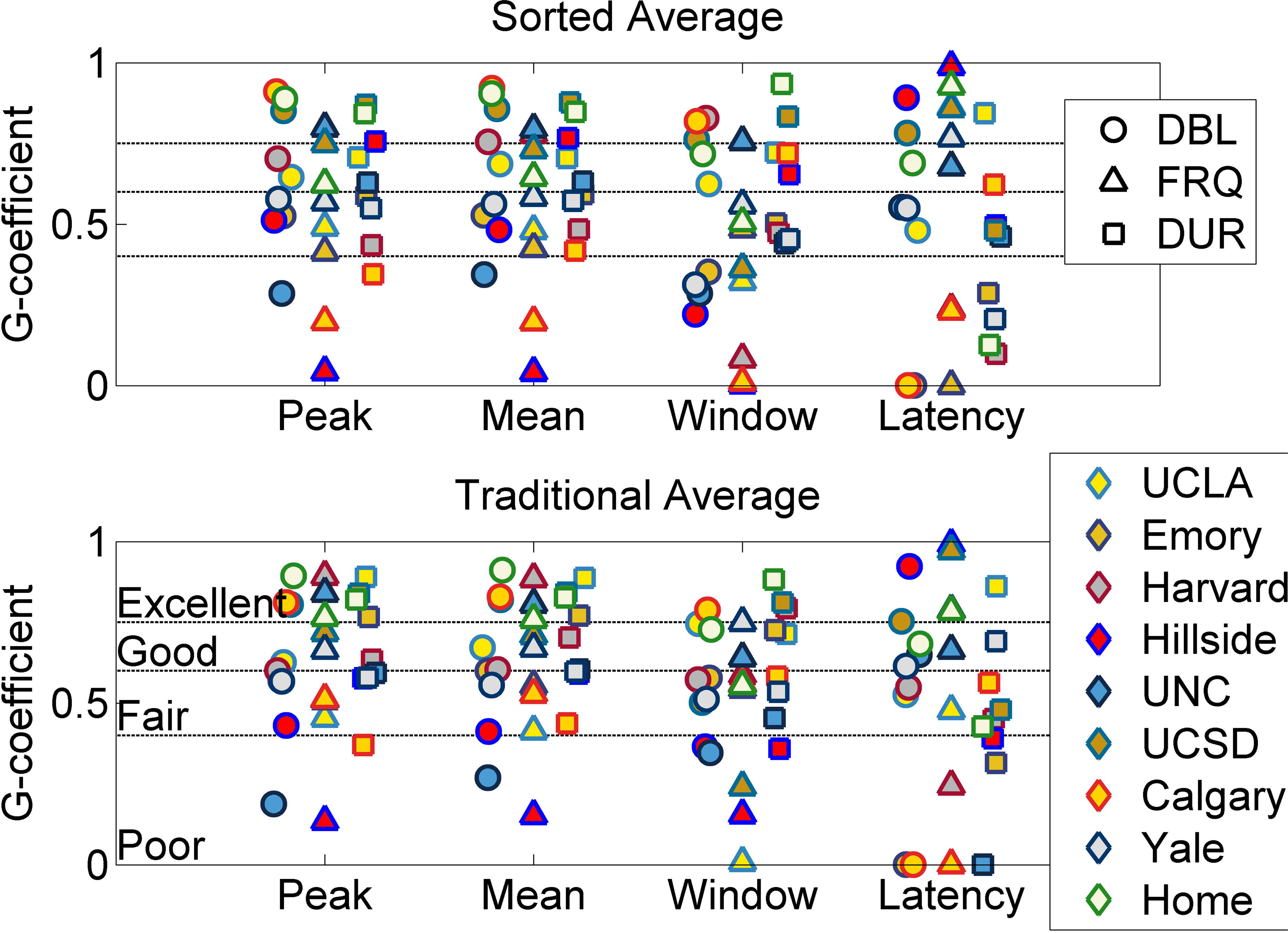
G-coefficients for the single-facet (test occasion) generalizability sub-studies calculated separately for each NAPLS geographic site and a 9^th^ “home” site G-study for electrode Fz based on either sorted averaging (top) or traditional averaging (bottom) event-related potential calculation. In all cases, measurement approaches are plotted along the x-axis separately for double-deviant (DBL, circles), frequency-deviant (FRQ, triangles), and duration-deviant (DUR, squares) mismatch negativity.

In the single-facet models, the DUR measures tend to have the smallest range, while the FRQ measures have the widest range of G-coefficients across measurement and averaging methods at electrode Fz. Of the 4 measures, the latency G-coefficient range is clearly the largest, with G-coefficients approaching the extremes. G-coefficients for each electrode, deviant type, reference, averaging approach, and measure calculated separately at each of the 8 sites (and a 9^th^ set for “home” site) are included in Table S2.

There were 420 (54.69%) home site G-coefficients greater than or equal to 0.6, which was more than (i) seven of eight NAPLS sites and (ii) all other pairs of subsequent, ordered visits. The Yale site had 429 (55.86%) G-coefficients that were good or better and the next closest pair of visits was the 6^th^ ordered site (visits 12 and 13) with 287 (37.37%). The home site G-coefficient was maximal in 167/768 (21.74%) comparisons with the 8 NAPLS sites and 200/768 (26.04%) comparisons with the 7 pairs of subsequent visits. Based on the 95% confidence intervals for these proportions, there is significant evidence that the home site is greater than all the other NAPLS sites more often than would be expected by chance (CI: 20.26– 23.23%, chance: 11.11%), and the home site is the greatest of any pair of ordered visits more often than would be expected by chance (CI: 24.46–27.63%, chance: 12.5%).

### Averaging Method and Reference Electrode Comparisons

Comparisons of the sorted and traditional averaging method, using mastoid referenced data, indicated that there was no difference between the methods when controlling for deviant type (CMH statistic = 2.22, df = 2, p = 0.329) or measure (CMH statistic = 2.04, df = 2, p = 0.361). This is consistent with G-coefficient category counts in Table 5.

**Table 5.**
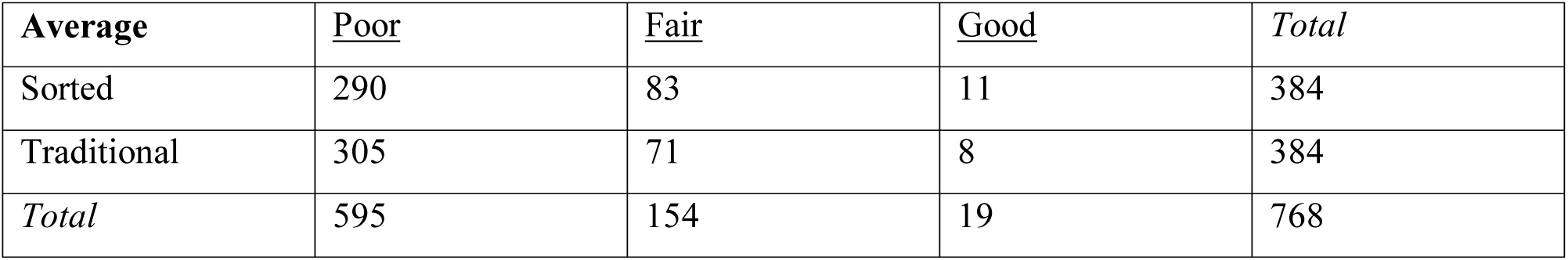
Frequency of G-coefficients by Averaging Method

Comparisons of the different reference electrodes indicated that there was a strong association between reference choice and G-coefficient categorization when adjusting for either deviant type (CMH statistic = 98.81, df = 2, p < 0.0001) or measure (CMH statistic = 93.81, df = 2, p < 0.0001). As seen in Table 6, the majority (∼94%) of nose referenced data G-coefficients were poor while only a small proportion (∼6%) were fair. The mastoid referenced data G-coefficients were fair or better in more than 22% of the cases.

**Table 6.**
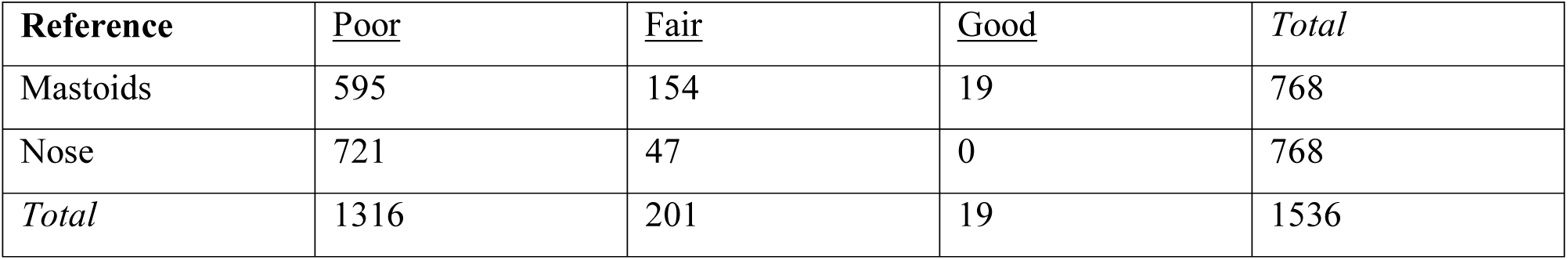
Frequency of G-coefficients by Reference Electrode

Given the relatively low frequency of G-coefficients above .6 (good), re-classification into two categories (above or below 0.4, the cut point for poor vs. fair) allowed us to assess the homogeneity of odds ratios across different strata (deviant or measure) using the Breslow-Day Test [5]. In both averaging method models, these tests were non-significant (deviant χ^2^ = 2.55, df = 2, p = 0.279; measure χ^2^ = 0.022, df = 2, p = 0.989). However, in both reference electrode models, there was significant statistical evidence of heterogeneous odds ratios (deviant χ^2^ = 10.93, df = 2, p = 0.0042; measure χ^2^ = 34.79, df = 2, p = <0.0001). When broken down by deviant type (Table S3), there was no difference in the odds of a fair or better G-coefficient for FRQ (95% CI: 0.62-3.50), 6.15 times greater odds of a fair or better G-coefficient for DUR (95% CI: 3.96-9.56), and 8.75 times greater odds of a fair or better G-coefficient for DBL (95% CI: 3.38-22.62) when data were mastoid referenced. When broken down by measure (Table S3), the odds of a fair or better G-coefficient were 8 times greater for the window measure (95% CI: 3.07-21.07), 5.67 times greater for the peak measure (95% CI: 3.22-9.99), 5.65 times greater for the mean measure (95% CI: 3.16-10.07), and 1.04 times worse for the latency measure (95% CI: 1.01-1.07) when data were mastoid referenced. On the whole, these results indicate that with the exception of the latency measure, mastoid referenced data always had greater odds of higher G-coefficients, and in particular the DBL deviant and window measures exhibited the strongest effects.

Alternatively, when ignoring these ICC categories and instead determining the proportion of G-coeffecients calculated using sorted average data that were greater than corresponding G-coefficients calculate using traditional averaging, 213 of 384 G-coefficients (55.47%) were greater. Based on the 95% confidence interval for this proportion (54.44 – 56.5%), there is evidence that the sorted averaging approach produces significantly more generalizable measures than traditional averaging (α = .05 level, two-tailed). The proportion of G-coefficients calculated using a mastoid reference that were greater than those using a nose reference was 74.74% (574 of 768 G-coefficients), indicating that the mastoid reference generally produced more reliable results. This effect was also statistically significant based on the 95% confidence interval (74.15 – 75.33%). To verify that the nose reference was not yielding poor reliability due to the order of operations (i.e., re-referencing the the nose electrode occurred after all pre-processing and artifact rejection), the order of operations was reversed in a parallel processing pipeline such that nose referencing occurred before all interpolation and artifact rejection procedures. However, the proportion of G-coefficients calculated with an initial nose reference that were greater than those using an average mastoid re-reference was only 15.23% (117 of 768 G-coefficients), indicating that order of operations does not explain poorer generalizability in the nose-referenced MMN scores.

## Discussion

The main purpose of this generalizability study was to quantify variance components and associated G-coefficients representing the single site, single session reliability of MMN measured with different methods of referencing, averaging, and scoring EEG data. Across these different cases, error variance was typically the largest, followed by either person or person by site variance components, depending on the type of response being measured (i.e., amplitude vs latency). All other variance components accounted for small proportions of variance in the study data. This pattern was consistent across delete-1 jackknife estimates of the variance components, with greatest variability in the person, person by site, and error variance components. While the range of the G-coefficients in these delete-1 models was non-trivial (∼0.2 to 0.6), this corresponds to the presence of a large person by site interaction variance component. Specifically, the amount of site variance depends on the particular person (alternatively, the person variance depends on the level of site), such that excluding a particular site, subject, or combination of site and subject might shrink the P x S interaction term and improve the G-coefficient.

Comparisons of MMN reliability using average mastoid versus nose references revealed that mastoid referenced data were more reliable in the majority (∼75%) of electrodes, deviants, measurements, and averaging approaches. Many test-retest reliability studies of MMN have used nose referenced data, most likely to show that the MMN component reverses polarity and that the associated scalp component is the MMN and not the N2b ERP component. Based on the findings from the current study, nose referencing appears to quite clearly increase relative error variance. Therefore, reports that MMN suffers from low or poor test-retest reliability that were based on a nose reference should be qualified as limited to MMN measures calculated using this particular reference.

The comparisons of reliability of MMN scores using two different averaging methods had mixed results. The sorted averaging approach, which removed an additional 6% of trials on average, only improved reliability estimates compared to a traditional averaging approach in slightly more than half (∼55%) of electrodes, deviants, and measurements. However, when limiting the focus to fronto-central electrodes and amplitude measures typically used in MMN group analyses, the difference in percentage of variance attributed to persons was less than 1.5%, on average, for these two averaging approaches. It is posible that the benefit of the sorted averaging algorithm is limited to electrodes where the signal is smaller, but this benefit seems to be very small, especially when one considers that the computation time and single trial implementation of the sorted averaging algorithm may be prohibitive for many EEG researchers.

Two previous studies reported excellent reliability (Fz ICCs > 0.85) using a long duration deviant similar to that used in the present study based on a window measurement (135-205 ms) from nose-referenced data [23, 24]. These high reliability coefficients were based on either 10 patients with schizophrenia tested twice, 18 months apart [23] or 163 patients and 58 comparison controls tested after 1 year [24]. Notably, the studies by Light et al used a much longer recording session that was terminated after a minimum of 225 artifact free deviant trials were obtained for each subject at run time and usually resulting in >250 trials following post-acquisition artifact correction procedures. While the corresponding traditionally averaged, mastoid referenced, window measure G-coefficient was smaller in the present study (Fz G = 0.492), the reduced “home” site model was almost identical (Fz G_home_ = 0.883). Another previous long duration deviant MMN reliability study of 19 healthy subjects tested twice, 7 to 56 days apart, had good reliability (Fz ICC = 0.66) using a left earlobe reference and similar window (50-200ms) measurement [18]. Lew et al. [22] also reported good reliability (Cz ICC = 0.6) in data from 19 healthy subjects tested twice, 2 to 60 days apart, using a frequency deviant and nose reference. However, unlike the previous two studies and the current G-study, the tones were part of an active auditory oddball attention task.

There are several limitations to the current study that should be carefully considered. First, estimates of variance components can be fairly unstable when the number of observations is small, and the estimates may have been impacted by having only 8 subjects studied on only two test occassions at each site. While our exact design is unlikely to be replicated in future reliability studies, focusing on a larger sample size and more repeat test occassions would yield more stable variance components estimates and potentially more informative decision studies. Second, the particular design employed here could have also introduced unintended psychological and/or physiological effects on the ERP measures. The participants completed the same EEG task 16 times, and 14 of these 16 test occasions involved some long travel times, which could have contributed to boredom, sleepiness, jetlag, and/or stress. This constrasts with previous MMN reliability studies that involve typically two, but at most four [11] or five [29], repeated assessments at one lab site. Finally, the geographic layout of the sites and administrative burden of organizing the study required a fixed travel loop for all subjects, and pseudo-randomization of order was achieved by having one subject start at each site. For example, all subjects visited UCSD after UCLA except for the subject who started at UCSD.

While order effects were not anticipated, they cannot be quantified in the current design and may have contributed to the variability of the within site reliability coefficients and the large Person X Site variance components in the fully crossed, two facet models. Despite these limitations, MMN measures have equal or greater reliability than task-based fMRI measures from this same cohort [15, 16].

In conclusion, the current study demonstrates the feasibility of a multisite, EEG studies of mismatch negativity using the same software and hardware. Grand average ERP waveforms were consistent across all NAPLS sites and test occasions (Figure 1) despite the small sample size and high number of repeated assessments conducted on each subject over a relatively brief assessment period. Moreover, when only the first two test occasions were considered from each traveling subject’s “home” site, reliability was good or better (i.e., G-coefficient or ICC > 0.6) in a large proportion of MMN measurements across electrodes. This home site design closely matches the main NAPLS design, in which subjects participate at a single site. Given the highly variable but generally poor reliability of latency measures, MMN amplitude assessed using a fixed latency, mean amplitude (e.g., “window” measure in this study) may be the most generalizable measure for multisite investigations of MMN in low prevalance patient groups, such as CHR individuals.

## Acknowledgements

This work was supported by grants from National Institute of Mental Health (U01MH081902 to TDC, P50 MH066286 to CEB, U01MH081988 to EFW, U01MH076989 to DHM, U01MH081944 to KSC, U01MH081984 to JA, U01MH082004 to DOP, U01MH081857 to BAC, U01MH081928 to LJS, U01MH082022 to SW).

## Disclaimer

The content is solely the responsibility of the authors and does not necessarily represent the official views of the National Institutes of Health or the Department of Veterans Affairs.

Drs Hamilton, Duncan, Johannesen, Light, Niznikiewicz, and Mathalon are employees of the US government.

