## Supplementary figures and images for "Reliability of Mismatch Negativity Event-Related Potentials in a Multisite, Traveling Subjects Study"

### Supplemental Figure 1

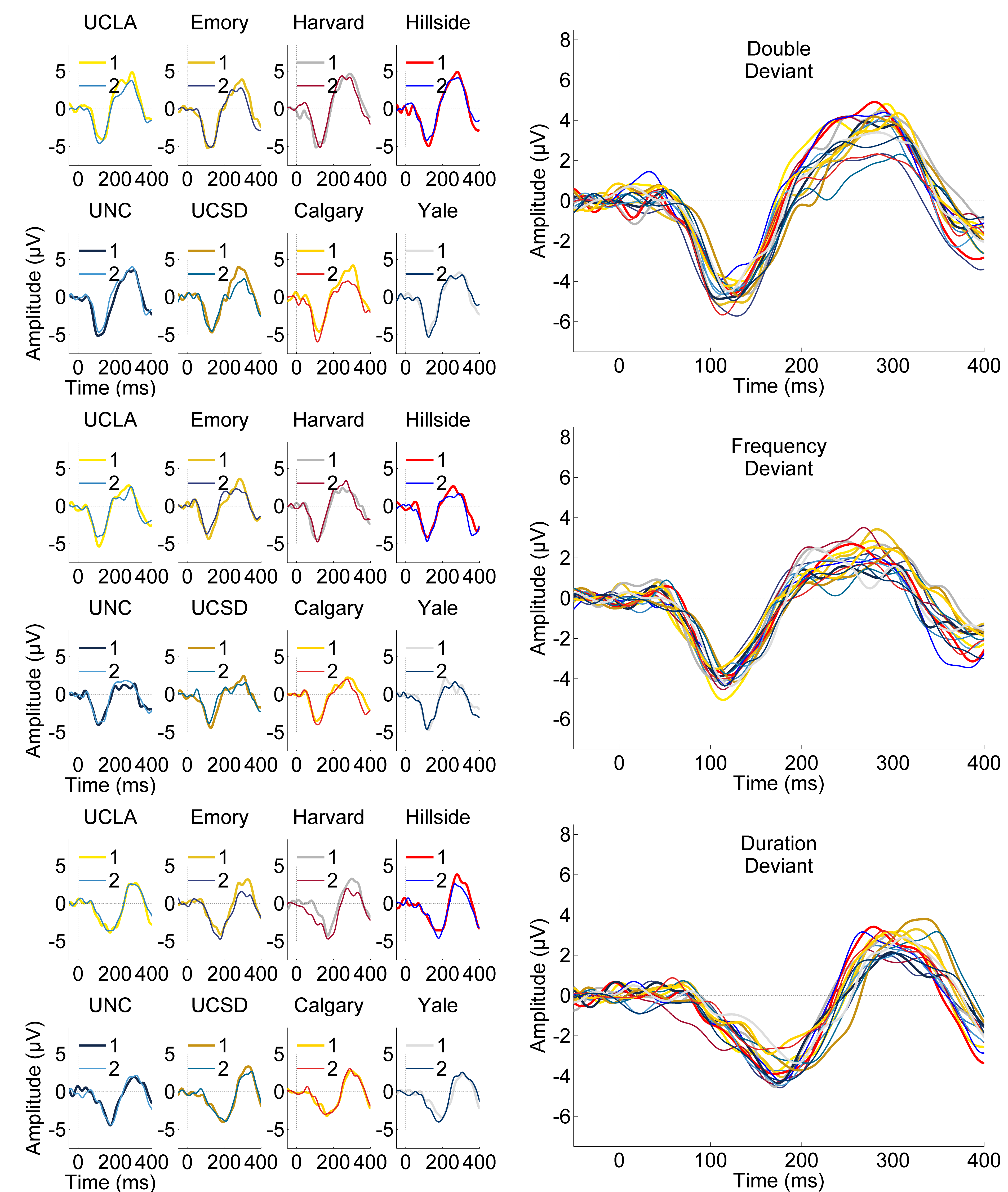

### Supplemental Figure 2

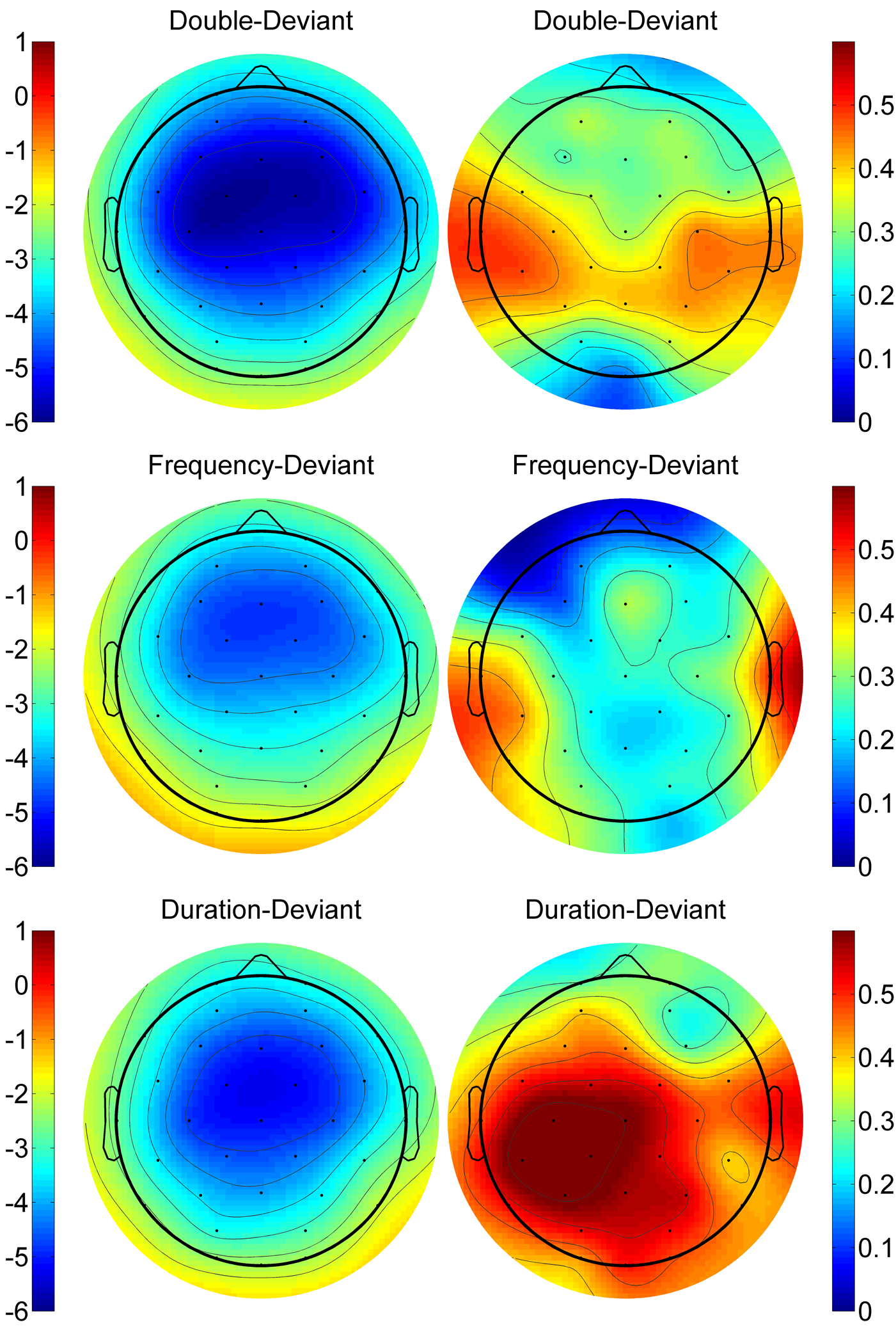

### Supplemental Figure 3

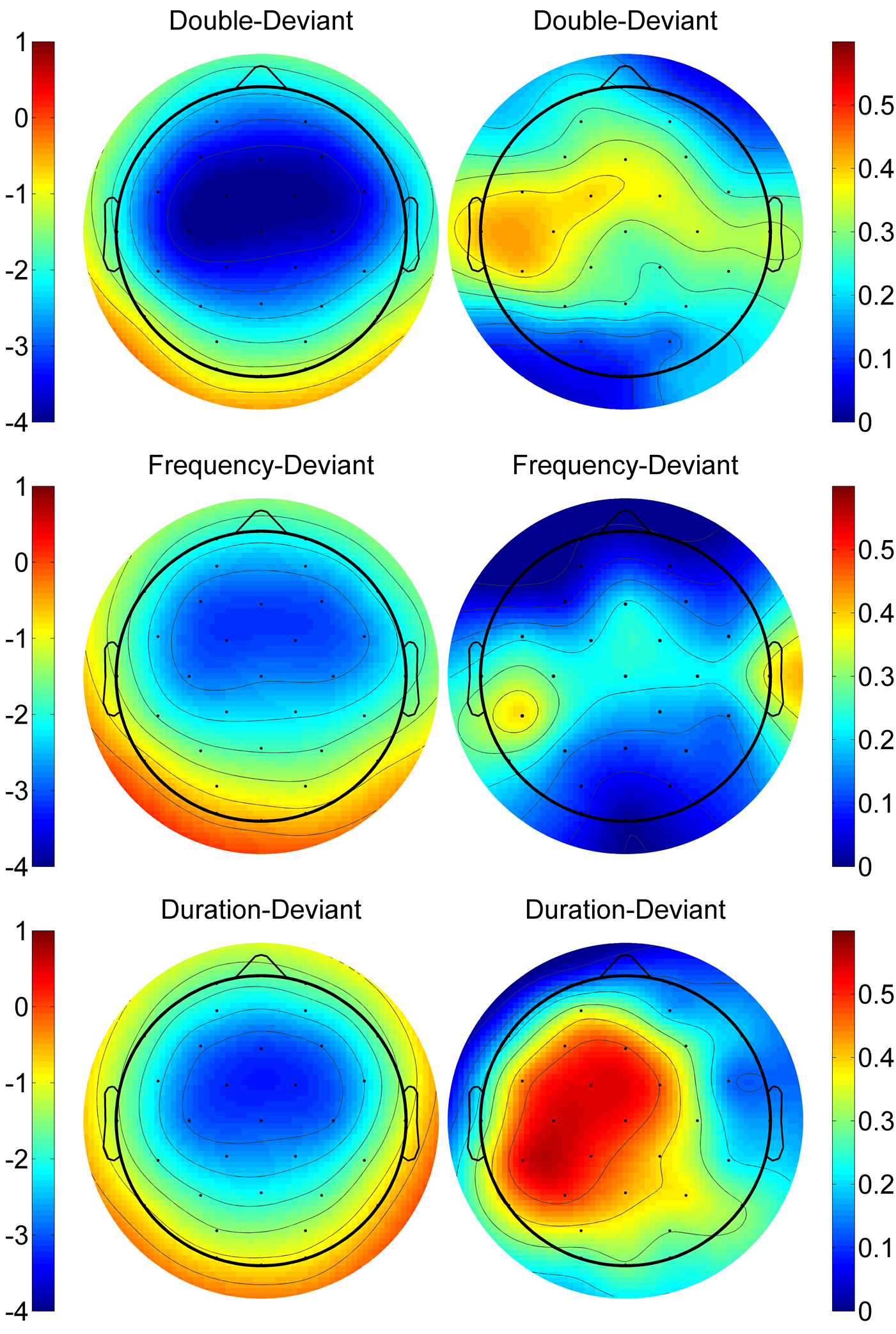

### Supplemental Figure 4

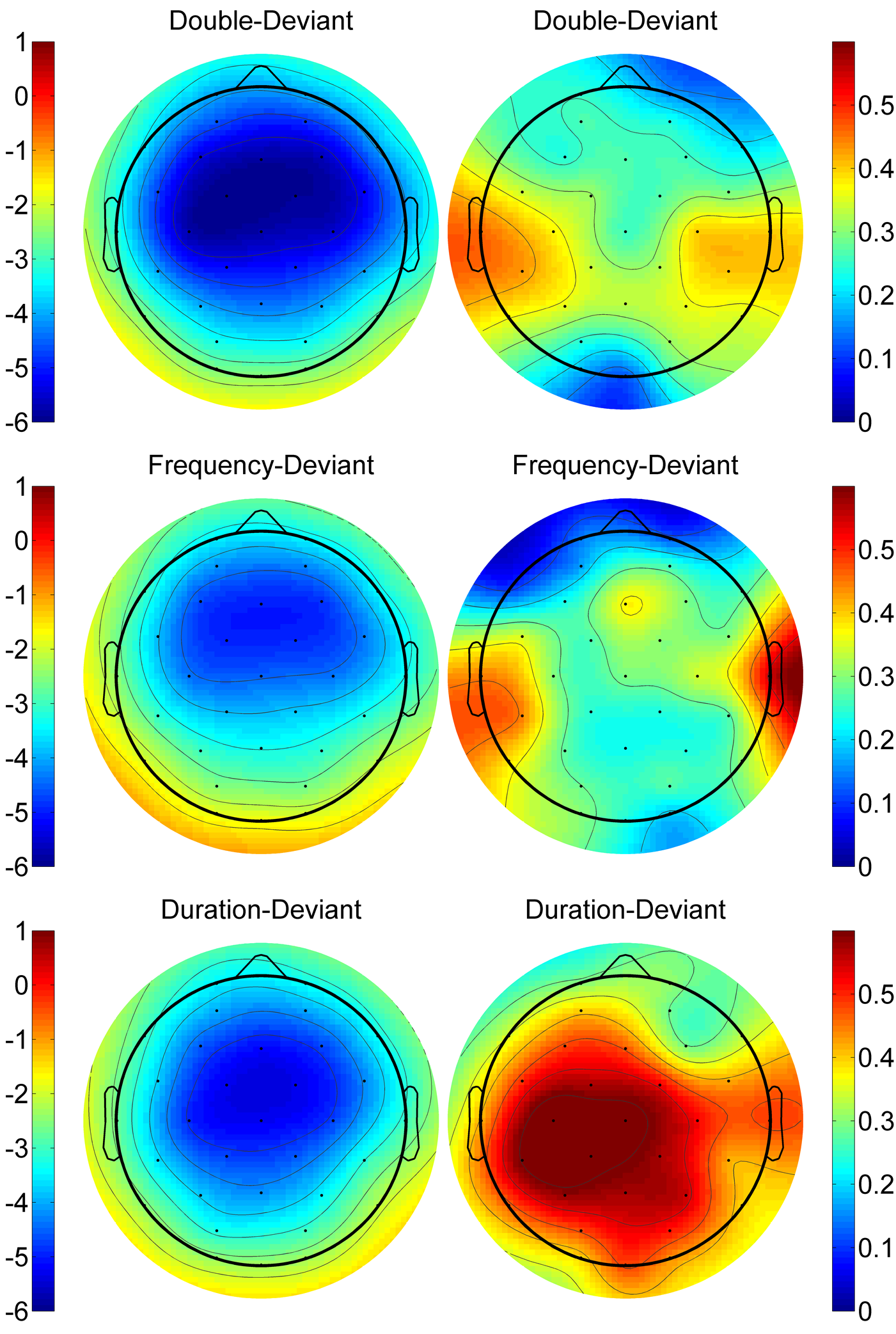
